# Resting heart rate and psychopathy: Findings from the Add Health Survey

**DOI:** 10.1101/205005

**Authors:** Nicholas Kavish, Q. John Fu, Michael G. Vaughn, Zhengmin Qian, Brian B. Boutwell

## Abstract

Despite the prior linkages of low resting heart rate to antisocial behavior broadly defined, less work has been done examining possible associations between heart rate to psychopathic traits. The small body of research on the topic that has been conducted so far seems to suggest an inverse relationship between the two constructs. A smaller number of studies have found the opposite result, however, and some of the previous studies have been limited by small sample sizes and unrepresentative samples. The current study attempts to help clarify the relationship between resting heart rate and psychopathic traits in a large, nationally representative sample (analytical *N* ranged from 14,173-14,220) using an alternative measure of psychopathic traits that is less focused on antisocial processes, and rooted in personality traits. No significant relationship between heart rate and psychopathic traits, or heart rate and a measure of cold heartedness, was found after controlling for age, sex, and race. Implications of the findings, study limitations, and directions for future research are discussed.

## Resting heart rate and psychopathy: Findings from the Add Health Survey

The past few decades have witnessed a resurgence of criminological research exploring potential biological predictors of antisocial and criminal behavior (Raine, 2002, 2015; Raine, Venables, & Williams, 1995; Sijtsema et al., 2010). One factor in particular, low autonomic arousal has emerged as a robust predictor of antisocial behavior across the life course, ranging from childhood into adulthood (Lorber, 2004; Ortiz & Raine, 2004; Raine, 2002). Indeed, multiple meta-analyses, analyzing dozens of empirical studies, have repeatedly suggested that low autonomic arousal, measured via resting heart rate, is predictive of aggression, conduct problems, and violence in juveniles. Providing further support for the link between autonomic arousal and crime, Raine, Venables, and Williams (1995) in a longitudinal study of adolescents in the UK found that a higher resting heart rate at age 15 was a protective factor against criminal behavior at age 29.

Two possible explanations have been offered to explain the link between resting heart rate and antisocial behavior. First, fearlessness theory suggests that lower levels of arousal are biomarkers of lower levels of fear (Raine, 1993). Individuals with lower resting heart rates, the argument goes, experience diminished levels of fear and anxiety and are less conditionable to the aversive consequences of their riskier behavior (Lykken, 1995; Raine, 2002). Owing to these diminished states of relatively low fear and anxiety, a resulting predisposition toward engaging in antisocial behavior or violence becomes more pronounced (Raine, 1993; 2002). Several studies have found that low physiological arousal correlates strongly with temperamental features of behavioral disinhibition early in the life-course, thus providing convergent evidence to support the fearlessness perspective (Fowles, Kochanska, & Murray, 2000; Kagan, 1994; Kagan, Reznick, & Snidman, 1987; Scarpa, Raine, Venables, & Mednick, 1997).

The second explanation that might account for the inverse correlation between heart rate and antisocial behavior suggests that individuals with lower levels of resting arousal tend to score higher on measures of sensation seeking (Eysenck & Eysenck 1967; Eysenck, 1977; Raine, Reynolds, Venables, Mednick, & Farrington, 1998). The explanation posits that individuals with low resting heart rates experience their low arousal states as uncomfortable and are driven to seek stimulation or novelty in order to increase their arousal to a more comfortable homeostatic level. Thus, antisocial behavior and violence are explained as the efforts of chronically under-aroused individuals engaging in sensation seeking behavior (Horvath & Zuckerman, 1993). Recent research supports this hypothesis with findings that sensation seeking mediates the relationship between low resting heart rate and aggression (Portnoy et al., 2014).

Most importantly for the current study, the constructs of both fearlessness and sensation seeking have been linked to psychopathy (Hare, 1965; Patrick, 1994), a construct that is also a consistent predictor of antisocial behavior and crime (Coid, 1998; Hare, 1991). Psychopathy represents a constellation of personality traits and behaviors, generally consisting of callousness, grandiosity, manipulativeness, dishonesty and impulsivity, as well as overt forms of antisocial behavior. Psychopaths also tend to exhibit deficits in emotion, remorse, and attention (Fox, Jennings, & Farrington, 2015; Frick & White, 2008; Hare, 1991, Patrick, Fowles, & Krueger, 2009). Especially pertinent to the current study, psychopathic individuals have also been found to be high on sensation seeking (Hare, 1991; Zuckerman, Buchsbaum, & Murphy, 1980) and prior research has often attempted to explain their behavior as a consequence, at least in part, of diminished fear arousal (Birbaumer et al., 2005; Hare, 1965; Patrick, 1994).

The consistent link between low autonomic arousal and antisocial behavior, coupled with the overlap between psychopathy and antisocial behavior, suggests an additional possible pathway leading from individual differences in arousal to individual differences in antisocial outcomes. To the extent that individuals with low levels of arousal also evince traits consonant with psychopathy, this may help clarify some of the reasons why low resting heart rate correlates with criminal behavior.

To date, however, there has been significantly less research examining the relationship between heart rate and psychopathy than has examined heart rate and antisocial behavior generally (Kavish et al. 2017). While one meta-analysis found no relationship between resting heart rate and psychopathy (Lorber, 2004), a subsequent systematic review and meta-analysis found an overall inverse relationship using different inclusion/exclusion criteria (*d* = −.19; Portnoy & Farrington, 2015). The existing research that Portnoy and Farrington (2015) analyzed has some important limitations. The first drawback of the existing literature is that the majority of the studies available to be included in the meta-analysis seem to have measured psychopathy via the PCL/PCL-R and other closely related measures (e.g. Lobbestael et al., 2009; Ogloff & Wong, 1990; Raine et al., 2014).

While the PCL-R is generally considered the gold standard measure for assessing psychopathy, there is also a substantial body of research that assesses psychopathy from a dimensional personality perspective (e.g. De Vries & van Kampen, 2010; Kavish, Sellbom, & Anderson, In Press; Lee & Ashton, 2005; Miller et al., 2001; Miller & Lynam, 2003). To date, however, there is a lack of research focusing on the relationship between physiological variables and psychopathic personality features. Furthermore, Portnoy and Farrington (2015) were only able to analyze eight effect sizes for the relationship between resting heart rate and factors of psychopathy, with five effect sizes for the interpersonal/affective Factor 1 and only three for the lifestyle/antisocial Factor 2. Analysis of these few studies revealed no significant differences between heart rates relation to the two factors; however, in addition to the small number of studies analyzed, evaluation of only a two-factor structure of psychopathic traits may obscure some degree of variability, as some research has suggested a four-factor model of psychopathy (Interpersonal, Affective, Lifestyle, Antisocial; Hare, 2003; Vitacco, Neumann, & Jackson, 2005; Williams, Paulhaus, & Hare, 2007). Indeed, a recent study found a significant inverse correlation between resting heart rate and some of the affective features of psychopathy (callousness, unemotionality) in juveniles (Kavish et al., 2017), a relationship that could not be assumed from prior research due to the combination of affective and interpersonal traits. Caution is warranted, however, given that the study by Kavish et al. (2017) was conducted using a small, non-representative sample.

Another important point regarding the research analyzed in Portnoy and Farrington (2015) is that the existing literature from which the authors had to draw contains an overabundance of effect sizes drawn from incarcerated and institutionalized samples, along with a lack of large, representative samples. Portnoy and Farrington (2015) themselves point to the overrepresentation of child and juvenile samples and the theoretically inconsistent finding of no moderating effect of age as evidence of a need for more research with adult populations. Given these limitations in the existing research, and thus the limitations placed on what the authors of the meta-analysis had to work with, further research is needed to better elucidate the possible link between heart rate and psychopathy.

### Current Study

The association between heart rate and psychopathy is a relatively novel finding, about which not much is deeply understood at the current juncture. Despite being able to possibly shed light on the link between heart rate and criminal behavior, this line of research is in its infancy and prior evidence (Kavish et al., 2017) has relied mostly on small samples and overt measures of antisocial behavior. Given the growing recognition within psychology of the need for replication of findings (Open Science Collaboration, 2015), including with alternative measures and diverse samples, we believe more work is needed on this topic. The current study uses a nationally representative sample of American respondents and a measure of psychopathy that is relatively free of antisocial processes in order to further scrutinize the association of resting heart rate and psychopathic personality tendencies.

## Methods

### Participants

To expand the prior work in this area (some of which, as we mentioned above, relied on relatively small samples), we analyzed data from the National Longitudinal Study of Adolescent to Adult Health (Add Health; Udry, 2003). The Add Health is a widely used data source of scholars across academic disciplines. The nature of the data collection, including sampling strategy, measurement, and availability of the data have been repeatedly described elsewhere (Beaver et al., 2013; Harris, Halpern, Smolen, & Haberstick, 2006). Briefly, the Add Health^1^ is a longitudinal and school-based survey derived from a nationally representative sample of adolescents in grades 7-12 first interviewed in 1994-1995 (Harris, Halpern, Smolen, & Haberstick, 2006). There have been a total of 4 waves of data collected and made available to date. The fourth, and most recent, wave of the study was conducted in 2008 with original sample aged 24-34 and the response rate was 80.3% (15,701 of the original participants completed Wave 4). Questions at Wave 4 largely addressed employment history, behavior, personality traits, and lifetime contact with the criminal justice system (Harris et al., 2003), and also included a measure of resting heart rate. The current study draws its data from Wave 4 of the Add Health study. Among 15,701 participants, our analytical sample size ranged from 14,173 – 14,220 due to incomplete or missing data on study variables (e.g., age, sex, race, heart rate, psychopathy, or cold-heartedness).

### Measures

#### Physiological arousal

Physiological arousal data for the current study is drawn from the cardiovascular measurements of Wave 4. In Wave 4, trained and certified field interviewers used factory calibrated Microlife BP3MC1-PC-IB oscillometric blood pressure monitors (MicroLife USA, Inc.; Dunedin, FL) to measure pulse rates in beats per minute. Measurements were taken 3 times at 30 second intervals. Participants were seated during the measurements and the 2^nd^ and 3^rd^ readings were averaged (Entzel et al., 2009). In line with prior research (Anselmino, Öhrvik, & Rydén, 2010; Carnethon et al., 2008; Maliphant, Watkins, & Davies, 2003; Murray et al., 2016; Sandset et al., 2014), the heart rate measure was divided in quartiles (i.e. 1^st^ quartile=66 or lower, 2^nd^ quartile=67-73, 3^rd^ quartile=74-82, and 4^th^ quartile=83 or higher).

#### Psychopathic personality styles

The Add Health did not include a specific measure of psychopathic traits. However, in Wave 4, respondents were asked questions which assessed their personality traits based on the Five Factor Model (FFM) of personality as well as their levels of self-control and self-regulation. The current study uses a psychopathic personality traits inventory (for further details on the development of this scale, see Beaver, Barnes, May, & Schwartz, 2011; Beaver, Rowland, Schwartz, & Nedelec, 2011) that was developed based on research suggesting that certain FFM items can be used to create a continuous measure of psychopathic personality traits (Derefinko & Lynam, 2007; Gudonis, Miller, Miller, & Lynam, 2008; Lynam, 2002; Lynam et al., 2005; Lynam & Derefinko, 2006; Lynam & Widiger, 2001; Miller & Lynam, 2003; Miller, Lynam, Widiger, & Leukefeld, 2001; but see Miller et al., 2011 and Samuel & Widiger, 2008). The measure’s 23 items, scored on a 5 item Likert scale (1 = Strongly Agree - 5 = Strongly Disagree), assess the respondents’ ability to sympathize, feel other’s emotions, and whether or not the respondent cares about others. Higher scores on this scale represent more psychopathic personality traits (α = .80) and it should be noted that this scale has previously been used in multiple other studies (Beaver, Barnes, et al., 2011; Beaver, Boutwell, et al., 2015; Beaver, Rowland, et al., 2011). The items for this scale are included in Appendix A.

#### Cold-heartedness

The Add Health was also designed without the inclusion of an explicit measure of callous and unemotional traits, but again did include items assessing FFM personality traits. Therefore, the current study utilized a Cold-heartedness scale comprised of 8 items in the survey. The same items were used in a callous-unemotional scale created by Markowitz, Ryan, and Marsh (2015). The original scale contained 10 items; however, we elected to use only the 8 items that, in our view, most closely mapped onto the ICU (Markowitz, Ryan, & Marsh, 2015). Higher scores on this scale represent more cold-heartedness traits. The items used in this scale are included in Appendix B.

#### Control variables

The current study used participants’ ages at Wave 4 and treated them as a continuous variable. We also controlled for race, which was coded as White, Black, or Other and White was used as the reference. Finally, sex was included and was coded as either Male or Female and Male is the reference.

### Statistical Analysis

The primary independent variable for the current study was the quartile of heart rate. In order to assess the relationship between physiological arousal and psychopathic personality, we utilized a straightforward analytical approach. Total psychopathy and cold-heartedness scores were calculated by summing the responses to each item on its respective scale. Then, we examined the relationship between the heart rate measure and psychopathy using ordinary least squares (OLS) regression and controlling for age, gender and race variables. We further examined the extent to which an affective personality feature was predicted by arousal by testing the relationship between heart rate and the cold-heartedness scale. The weighted means, regression coefficients and standard errors were reported. Statistical significance was determined by p-value < 0.05. The design variables and sampling weights were incorporated in the analysis using SAS survey procedures (SAS Institute Inc. 2015).

## Results

Table 1 depicts the mean values of the FFM psychopathy and cold-heartedness scales. As we mentioned, higher scores represent a more psychopathic and cold-hearted character.

**Table 1.** Descriptive characteristics of the sample

|  | Mean | SE | Minimum | Maximum | *n* |
| --- | --- | --- | --- | --- | --- |
| Psychopathy | 66.85 | 0.09 | 41.00 | 98.00 | 14,800 |
| Cold Heartedness | 21.81 | 0.06 | 8.00 | 40.00 | 14,800 |
| Heart Rate | 75.04 | 0.55 | 39.00 | 200.00 | 14,800 |
| Age | 28.49 | 0.12 | 24.00 | 34.00 | 14,800 |
| Male % | 50.68 | 0.62 |  |  | 14,800 |
| Race |  |  |  |  | 14,774 |
| White | 79.18 | 2.39 |  |  |  |
| African-American | 16.67 | 2.14 |  |  |  |
| Other | 4.16 | 0.85 |  |  |  |

Note: SE: standard errors.

Table 2 presents the relationship between psychopathy and resting heart rate as well as the cold-heartedness measure and resting heart rate using linear regression. Resting heart rate was associated with psychopathy and cold-heartedness (at the bivariate level). Specifically, the estimated psychopathy scores were significantly greater in the first quartile of resting heart rate (e.g. heart rate<66/minute) than in the fourth quartile of resting heart rate (i.e. heart rate>81/minutes). The estimated cold-heartedness scores were higher in the first and second quartile of resting heart rate (e.g. heart rate<66/minute and 66<heart rate<73.5/minute) than in the fourth quartile of resting heart rate (i.e. heart rate>81/minutes), respectively. While it is common to examine the effects of resting heart rate across quartiles, we also analyzed our models using resting heart rate as a continuous variable. In general, the result remained unchanged. In particular, some evidence of a bivariate association between HR and coldheartedness emerged. Nonetheless, HR failed to evince a significant effect in the multiple regression model for either psychopathy or cold-heartedness.

**Table 2.** Bivariate analysis for psychopathic personality and cold heartedness

|  | Model 1: Psychopathy |  | Model 2: Cold Heartedness |  |
| --- | --- | --- | --- | --- |
|  | *b* | SE | *b* | SE |
| 1<sup>st</sup> quartile of heart rate | 0.40* | 0.20 | 0.55** | 0.14 |
| 2<sup>nd</sup> quartile of heart rate | 0.32 | 0.22 | 0.44** | 0.12 |
| 3<sup>rd</sup> quartile of heart rate | -0.05 | 0.19 | 0.05 | 0.12 |
| 4<sup>th</sup> quartile of heart rate | Reference |  | Reference |  |
| *n* | 14,173 |  | 14,220 |  |

Note: *b*: slope; SE: standard error; \* Significant at $p < 0.05$ ; \*\* Significant at $p < 0.01$ .

After controlling for sex, age and race, neither of the relationships between resting heart rate and psychopathy or the association between resting heart rate and cold-heartedness was significant. Table 3 contains the results of the two models that include the additional covariates. Our analyses, contrary to predictions and earlier research, found no significant relationship between any quartile of resting heart rate and scores on the psychopathy measure after adjustment for sex, age, and race. Similarly, no relationship was found between resting heart rate and scores on the cold-heartedness scale following these adjustments. Each of the demographic variables was significantly associated with both psychopathy and cold-heartedness personality traits. Increased age and being female were significantly associated with lower psychopathy and cold-heartedness scores. African Americans and those identifying as Other were associated with greater psychopathy and cold-heartedness scores than Whites. Implications of the results and study limitations are discussed.

**Table 3.** OLS regression analysis for psychopathic personality and cold heartedness

|  | Model 1: Psychopathy |  | Model 2: Cold Heartedness |  |
| --- | --- | --- | --- | --- |
|  | *b* | SE | *b* | SE |
| 1<sup>st</sup> quartile of heart rate | -0.29 | 0.18 | 0.01 | 0.12 |
| 2<sup>nd</sup> quartile of heart rate | -0.04 | 0.20 | 0.16 | 0.11 |
| 3<sup>rd</sup> quartile of heart rate | -0.14 | 0.17 | -0.04 | 0.10 |
| 4<sup>th</sup> quartile of heart rate | Reference |  | Reference |  |
| Age | -0.12** | 0.04 | -0.05* | 0.03 |
| Female | -3.58** | 0.14 | -2.71** | 0.08 |
| Race |  |  |  |  |
| Black | 0.70** | 0.19 | 0.89** | 0.13 |
| Other | 0.09 | 0.34 | 0.45** | 0.17 |
| White | Reference |  | Reference |  |
| *n* | 14,173 |  | 14,220 |  |

*b*: slope; Note: SE: standard error; \* Significant at $p < 0.05$ . \*\* Significant at $p < 0.01$ .

## Discussion

The current study sought to replicate and extend the existing body of research on the relationship between low resting heart rate and psychopathy by using a measure that is relatively free of antisocial behavior. To date, research has been mixed on the effects of resting heart rate on psychopathic traits (Lorber, 2004; Portnoy & Farrington, 2015). Most recently, Kavish et al. (2017) using a small convenience sample uncovered a significant effect of heart rate on scores on the Inventory of Callous Unemotional Traits (ICU) as well as the Youth Psychopathy Inventory (YPI). Given the small circumscribed sample, it remains an open question whether similar findings will emerge using alternative and more generalizable samples.

We sought to expand the analysis of Kavish et al. (2017) by examining other personality constructs—cold-heartedness—in order to try and further unpack the effect of heart rate on personality constructs that increase the risk of antisocial and criminal behavior. Consistent with previous work, we found that low resting heart rate was associated with psychopathic personality traits in the bivariate analyses. However, when sex, age and race were taken into account, our results failed to support the conclusions of previous work by revealing no significant relationship between heart rate and psychopathic personality features. The emergence of age and sex in the models, in particular, are not surprising given previous research which has found resting heart rate differs with age (Shinebourne, 1974 as cited in Raine, Venables, & Mednick, 1997) and by sex, even explaining as much as 17% of the gender gap in crime in one sample (Choy, Raine, Venables, & Farrington, 2017).

It is important to consider the limitations inherent in the current study. First, and perhaps foremost, the measure used to assess psychopathic personality features is based on items drawn from FFM personality items, and not from typical clinical measures of psychopathy. Prior research has utilized this measure before, and the constituent items appear to possess adequate validity (Beaver, Barnes, May, & Schwartz, 2011; Beaver, Rowland, Schwartz, & Nedelec, 2011), yet the extent to which the measure captures the same variation in psychopathic tendencies as other classic measures of psychopathy remains an open question. Furthermore, when Portnoy and Farrington (2015) unpacked psychopathy into Factors 1 and 2 in their meta-analysis, they only found a significant relationship between resting heart rate and Factor 2 (the lifestyle and antisocial behavior component). Combined with the firmly established link between heart rate and antisocial behavior (Ortiz & Raine, 2004; Portnoy & Farrington, 2015), this may suggest that heart rate is more likely to be associated with measures of psychopathy that include overt antisocial behavior. Similarly, the cold-heartedness scale was constructed from questions in the Wave 4 survey, as formal measures of callous and unemotional traits were not utilized in the Add Health data collection. The cold-heartedness scale has not been validated by other research and is in need of further study before strong conclusions can be drawn. Finally, the analysis was cross-sectional, with both measures assessed at Wave 4. This is perhaps less relevant, given the lack of significant findings. Nonetheless, it warrants mentioning as a longitudinal analysis would have been preferable, and should be conducted when additional waves of the Add Health become available.

Despite its drawbacks, the current study still provides an important contribution to the literature. There has been a great deal of concern recently regarding the methodological rigor and reproducibility of research across the field of psychology, and across science broadly (Benjamin et al., 2017; Crandall & Sherman, 2016), following a widely publicized failure to replicate a large number of famous psychological studies (Open Science Collaboration, 2015). These conversations have been a reminder of the importance of replication of past research: both conceptually and with alternative samples and measures in order to assess the generalizability of a given finding. In the current study, we utilized a large, national sample of adults, as well as an alternative method of assessing psychopathic traits to further investigate findings from previous research. Although the scale used in this study may not capture the antisocial behavior components assessed in traditional measures of psychopathy, it does provide an opportunity to examine the relationship between heart rate and the psychopathy construct from a FFM personality perspective. Ultimately, the current study emphasizes the necessity for more research into the relationship between resting heart rate and psychopathy using large, representative samples and alternative measures along with rigorous controls.

## Appendix A

Items included in the Psychopathic Personality Styles scale

1. I sympathize with others’ feelings
2. I get angry easily*
3. I am not interested in other people’s problems*
4. I often forget to put things back in their proper place*
5. I am relaxed most of the time
6. I am not easily bothered by things
7. I rarely get irritated
8. I talk to a lot of different people at parties
9. I feel others’ emotions
10. I get upset easily*
11. I get stressed out easily*
12. I lose my temper*
13. I keep in the background*
14. I am not really interested in others*
15. I seldom feel blue
16. I don’t worry about things that have already happened
17. I keep my cool
18. I go out of my way to avoid having to deal with problems in my life*
19. When making a decision, I go with my “gut feeling” and don’t think much about the consequences of each alternative*
20. I live my life without much thought for the future*
21. Other people determine most of what I can and cannot do*
22. There are many things that interfere with what I want to do*
23. There is really no way I can solve the problems I have*

## Appendix B

Items included in the Cold-heartedness scale:

1. I sympathize with others’ feelings *
2. I worry about things*
3. I am not interested in other people’s problems
4. I am relaxed most of the time
5. I am not easily bothered by things
6. I feel others’ emotions*
7. I am not really interested in others
8. I keep my cool

## Footnotes

http://www.cpc.unc.edu/projects/addhealth/design

* denotes reverse scoring

* denotes reverse scoring

## References

Anselmino, M., Öhrvik, J., & Rydén, L. (2010). Resting heart rate in patients with stable coronary artery disease and diabetes: a report from the euro heart survey on diabetes and the heart. European Heart Journal, 31(24), 3040–3045.

Beaver, K. M., Barnes, J. C., May, J. S., & Schwartz, J. A. (2011). Psychopathic personality traits, genetic risk, and gene–environment correlations. Criminal Justice and Behavior, 38, 896–912.

Beaver, K. M., Boutwell, B. B., Barnes, J. C., Vaughn, M. G., & DeLisi, M. (2015). The Association Between Psychopathic Personality Traits and Criminal Justice Outcomes Results From a Nationally Representative Sample of Males and Females. Crime & Delinquency 63(6), 1–23.

Beaver, K. M., DeLisi, M., Wright, J. P., Boutwell, B. B., Barnes, J. C., & Vaughn, M. G. (2013). No evidence of racial discrimination in criminal justice processing: Results from the National Longitudinal Study of Adolescent Health. Personality and Individual Differences, 55(1), 29–34.

Beaver, K. M., Rowland, M. W., Schwartz, J. A., & Nedelec, J. L. (2011). The genetic origins of psychopathic personality traits in adult males and females: Results from an adoption–based study. Journal of Criminal Justice, 39, 426–432.

Benjamin, D. J., Berger, J. O., Johannesson, M., Nosek, B. A., Wagenmakers, E. J., Berk, R., … & Cesarini, D. (2017). Redefine statistical significance. Nature Human Behaviour.

Birbaumer, N., Veit, R., Lotze, M., Erb, M., Hermann, C., Grodd, W., & Flor, H. (2005). Deficient fear conditioning in psychopathy: a functional magnetic resonance imaging study. Archives of General Psychiatry, 62(7), 799–805.

Carnethon, M. R., Yan, L., Greenland, P., Garside, D. B., Dyer, A. R., Metzger, B., & Daviglus, M. L. (2008). Resting heart rate in middle age and diabetes development in older age. Diabetes Care, 31(2), 335–339.

Choy, O., Raine, A., Venables, P. H., & Farrington, D. P. (2017). Explaining the gender gap in crime: The role of heart rate. Criminology, 55(2), 465–487.

Coid, J. W. (1998). The management of dangerous psychopaths in prison. In T. Milolon, E. Simonsen, M. Birket–Smith, & R. D. Davis (Eds.), Psychopathy: Antisocial, criminal, and violent behavior (pp. 431–457). New York: Guilford.

Crandall, C. S., & Sherman, J. W. (2016). On the scientific superiority of conceptual replications for scientific progress. Journal of Experimental Social Psychology, 66, 93–99.

Derefinko, K. J., & Lynam, D. R. (2007). Using the FFM to conceptualize psychopathy: A test using a drug abusing sample. Journal of Personality Disorders, 21, 638–656.

De Vries, R. E., & van Kampen, D. (2010). The HEXACO and 5DPT models of personality: a comparison and their relationships with psychopathy, egoism, pretentiousness, immorality, and Machiavellianism. Journal of Personality Disorders, 24(2), 244–257.

Entzel, P., Whitsel, E. A., Richardson, A., Tabor, J., Hallquist, S., Hussey, J., …Harris, K. M. (2009). Add Health Wave IV documentation: Cardiovascular and anthropometric measures. Retrieved from http://www.cpc.unc.edu/projects/addhealth/data/guides/Wave%20IV%20cardiovascular%20and%20anthropometric%20documentation%20110209.pdf.

Eysenck, H. J. (1977). Crime and personality (3rd ed.). St. Albans: Paladin

Eysenck, H. J., & Eysenck, S. B. (1967). On the unitary nature of extraversion. Acta Psychologica, 26, 383–390.

Fowles, D. C., Kochanska, G., & Murray, K. (2000). Electrodermal activity and temperament in preschool children. Psychophysiology, 37, 777–787.

Fox, B. H., Jennings, W. G., & Farrington, D. P. (2015). Bringing psychopathy into developmental and life–course criminology theories and research. Journal of Criminal Justice, 43(4), 274–289.

Frick, P. J., & White, S. F. (2008). Research review: The importance of callous–unemotional traits for developmental models of aggressive and antisocial behavior. Journal of Child Psychology and Psychiatry, 49(4), 359–375.

Gudonis, L. C., Miller, D. J., Miller, J. D., & Lynam, D. R. (2008). Conceptualizing personality disorders from a general model of personality functioning: Antisocial personality disorder and the Five–Factor Model. Personality and Mental Health, 2, 249–264.

Hare, R. D. (1965). Psychopathy, fear arousal and anticipated pain. Psychological Reports, 16(2), 499–502.

Hare, R. D. (1991/2003). The Hare psychopathy checklist–Revised. Toronto, Ontario, Canada: Multi–Health Systems.

Harris, K. M., Florey, F., Tabor, J., Bearman, P. S., Jones, J., & Udry, J. R. (2003). The National Longitudinal Study of Adolescent Health: Research design. Retrieved from http://www.cpc.unc.edu/projects/addhealth/design

Harris, K. M., Halpern, C. T., Smolen, A., & Haberstick, B. C. (2006). The national longitudinal study of adolescent health (Add Health) twin data. Twin Research and Human Genetics, 9(06), 988–997.

Horvath, P., & Zuckerman, M. (1993). Sensation seeking, risk appraisal, and risky behavior. Personality and Individual Differences, 14(1), 41–52.

Kagan, J. (1994). Galen’s prophecy: Temperament in human nature. New York: Basic Books.

Kagan, J., Reznick, J. S., & Snidman, N. (1987). The physiology and psychology of behavioral inhibition in children. Child Development 58(6), 1459–1473.

Kavish, N., Sellbom, M., & Anderson, J. L. (In Press). Implications for the Measurement of Psychopathy in the DSM–5 Using the Computerized Adaptive Test of Personality Disorder (CAT–PD). Journal of Personality Assessment.

Kavish, N., Vaughn, M.G, Cho, E., Barth, A., Boutwell, B., Vaughn, S., Capin, P., Stillman, S., & Martinez, L. (2017) Physiological Arousal and Juvenile Psychopathy: Is Low Resting Heart Rate Associated With Affective Dimensions? Psychiatric Quarterly, 88(1) 103–114.

Lobbestael, J., Arntz, A., Cima, M., & Chakhssi, F. (2009). Effects of induced anger in patients with antisocial personality disorder. Psychological medicine, 39(4), 557–568.

Lorber, M. F. (2004). Psychophysiology of aggression, psychopathy, and conduct problems: a meta–analysis. Psychological Bulletin, 130(4), 531–552.

Lykken, D. T. (1995). The antisocial personalities. Hillsdale, NJ: Erlbaum.

Lynam, D. R. (2002). Psychopathy from the perspective of the Five Factor Model. In P. T. Costa & T. A. Widiger (Eds.), Personality disorders and the Five Factor Model of personality (2nd ed., pp. 325–350). Washington, DC: American Psychological Association.

Lynam, D. R., Caspi, A., Moffitt, T. E., Raine, A., Loeber, R., & Stouthamer–Loeber, M. (2005). Adolescent psychopathy and the Big Five: Results from two samples. Journal of Abnormal Child Psychology, 33, 431–443.

Lynam, D. R., & Derefinko, K. J. (2006). Psychopathy and personality. In C. J. Patrick (Ed.), Handbook of psychopathy (pp. 133–155). New York, NY: Guilford Press.

Lynam, D. R., & Widiger, T. A. (2001). Using the Five–Factor Model to represent the DSM–IV personality disorders: An expert consensus approach. Journal of Abnormal Psychology, 110, 401–412.

Maliphant, R., Watkins, C., & Davies, J. G. (2003). Disruptive behaviour in non–referred mainstream school children, aged seven to nine: A psychophysiological contribution. Educational Psychology, 23(4), 437–455.

Markowitz, A. J., Ryan, R. M., & Marsh, A. A. (2015). Neighborhood income and the expression of callous–unemotional traits. European Child & Adolescent Psychiatry, 24(9), 1103–1118.

Miller, J. D., Gaughan, E. T., Maples, J., & Price, J. (2011). A comparison of agreeableness scores from the Big Five Inventory and the NEO PI–R: Consequences for the study of narcissism and psychopathy. Assessment, 18(3), 335–339.

Miller, J. D., & Lynam, D. R. (2003). Psychopathy and the Five Factor Model of personality: A replication and extension. Journal of Personality Assessment, 81, 168–178.

Miller, J. D., Lynam, D. R., Widiger, T., & Leukefeld, C. (2001). Personality disorders as extreme variants of common personality dimensions: Can the Five Factor Model adequately represent psychopathy? Journal of Personality, 69, 253–276.

Murray, J., Hallal, P. C., Mielke, G. I., Raine, A., Wehrmeister, F. C., Anselmi, L., & Barros, F. C. (2016). Low resting heart rate is associated with violence in late adolescence: a prospective birth cohort study in Brazil. International Journal of Epidemiology, 45(2), 491–500.

Ogloff, J. R., & Wong, S. (1990). Electrodermal and cardiovascular evidence of a coping response in psychopaths. Criminal Justice and Behavior, 17(2), 231–245.

Open Science Collaboration. (2015). Estimating the reproducibility of psychological science. Science, 349. doi:10.1126/science.aac4716

Ortiz, J., & Raine, A. (2004). Heart rate level and antisocial behavior in children and adolescents: A meta–analysis. Journal of the American Academy of Child & Adolescent Psychiatry, 43(2), 154–162.

Patrick, C. J. (1994). Emotion and psychopathy: Startling new insights. Psychophysiology, 31(4), 319–330.

Patrick, C. J., Fowles, D. C., & Krueger, R. F. (2009). Triarchic conceptualization of psychopathy: Developmental origins of disinhibition, boldness, and meanness. Development and Psychopathology, 21(03), 913–938.

Portnoy, J., & Farrington, D. P. (2015). Resting heart rate and antisocial behavior: An updated systematic review and meta–analysis. Aggression and Violent Behavior, 22, 33–45.

Portnoy, J., Raine, A., Chen, F. R., Pardini, D., Loeber, R., & Jennings, J. R. (2014). Heart rate and antisocial behavior: the mediating role of impulsive sensation seeking. Criminology, 52(2), 292–311.

Raine, A. (1993). The psychopathology of crime: Criminal behavior as a clinical disorder. San Diego: Academic Press.

Raine, A. (2002). Biosocial studies of antisocial and violent behavior in children and adults: A review. Journal of Abnormal Child Psychology, 30(4), 311–326.

Raine, A. (2015). Low resting heart rate as an unequivocal risk factor for both the perpetration of and exposure to violence. JAMA Psychiatry, 72(10), 962–964.

Raine, A., Fung, A. L. C., Portnoy, J., Choy, O., & Spring, V. L. (2014). Low heart rate as a risk factor for child and adolescent proactive aggressive and impulsive psychopathic behavior. Aggressive Behavior, 40(4), 290–299.

Raine, A., Reynolds, C., Venables, P. H., Mednick, S. A., & Farrington, D. P. (1998). Fearlessness, stimulation–seeking, and large body size at age 3 years as early predispositions to childhood aggression at age 11 years. Archives of General Psychiatry, 55(8), 745–751.

Raine, A., Venables, P. H., & Mednick, S. A. (1997). Low resting heart rate at age 3 years predisposes to aggression at age 11 years: Evidence from the Mauritius Child Health Project. Journal of the American Academy of Child & Adolescent Psychiatry, 36(10), 1457–1464.

Raine, A., Venables, P. H., & Williams, M. (1995). High autonomic arousal and electrodermal orienting at age 15 years as protective factors against criminal behavior at age 29 years. The American Journal of Psychiatry, 152(11), 1595–1600.

Sandset, E. C., Berge, E., Kjeldsen, S. E., Julius, S., Holzhauer, B., Krarup, L. H., & Hua, T. A. (2014). Heart rate as a predictor of stroke in high–risk, hypertensive patients with previous stroke or transient ischemic attack. Journal of Stroke and Cerebrovascular Diseases, 23(10), 2814–2818.

Samuel, D. B., & Widiger, T. A. (2008). A meta–analytic review of the relationships between the five–factor model and DSM–IV–TR personality disorders: A facet level analysis. Clinical Psychology Review, 28(8), 1326–1342.

SAS Institute Inc. 2015. SAS/STAT^®^ 14.1 User’s Guide. Cary, NC: SAS Institute Inc.

Scarpa, A., Raine, A., Venables, P. H., & Mednick, S. A. (1997). Heart rate and skin conductance in behaviorally inhibited Mauritian children. Journal of Abnormal Psychology, 106, 182–190.

Shinebourne E. A. (1974). Growth and development of the cardiovascular system. In J. A. Davis & J. Dobbing (Eds.) Scientific Foundations of Pediatrics,(pp 206–214). Philadelphia, PA: Heinemann.

Sijtsema, J. J., Veenstra, R., Lindenberg, S., van Roon, A. M., Verhulst, F. C., Ormel, J., & Riese, H. (2010). Mediation of sensation seeking and behavioral inhibition on the relationship between heart rate and antisocial behavior: The TRAILS study. Journal of the American Academy of Child & Adolescent Psychiatry, 49(5), 493–502.

Udry, J. R. (2003). The National Longitudinal Study of Adolescent Health (Add Health), Waves I and II, 1994–1996; Wave III, 2001–2002 [Machine–readable data file and documentation]. Chapel Hill: Carolina Population Center, University of North Carolina at Chapel Hill.

Williams, K. M., Paulhus, D. L., & Hare, R. D. (2007). Capturing the four–factor structure of psychopathy in college students via self–report. Journal of Personality Assessment, 88(2), 205–219.

Zuckerman, M., Buchsbaum, M. S., & Murphy, D. L. (1980). Sensation seeking and its biological correlates. Psychological Bulletin, 88(1), 187–214.

